# Rethinking Molecular Beauty in the Deep Learning Era

**DOI:** 10.64898/2025.12.03.692079

**Authors:** Antoine Daina, Vincent Zoete

## Abstract

As (deep) generative chemistry rapidly enters the landscape of drug discovery, evaluating the models and their structural output relevance, tractability, and innovation potential becomes both a scientific and practical challenge. Existing metrics such as QED and SA Score, though widely adopted, are rooted in biased historical datasets and often fail to capture what medicinal chemists truly seek: compounds with meaningful pharmacological potential and realistic tractability.

In this work, we introduce the OiiSTER-map—a simple, intuitive, and interpretable two-dimensional heatmap that classifies molecules based on bioactivity-informed usuality and structural elaboration derived from molecular fingerprints. Unlike traditional filters, the OiiSTER-map helps identify not only the drug discovery chemical “sweet spot”. (Regular/Balanced), but also underappreciated territories such as Minimal/Unusual or Over-elaborated/Trivial regions, offering actionable insights into compound quality, relevance, and diversity.

We hope that a bioactivity-informed, structurally aware, and easily interpretable tool like the OiiSTER-map, employed in combination with other well implemented metrics, can be decisive to go beyond current limitations of assessing (deep) generative models and to ensure a more mechanistically relevant, nuanced and useful evaluation of the thus designed virtual molecules.

## Introduction

Today, it is acknowledged that the implementation of deep learning (DL) has benefited drug discovery. It is also widely anticipated that thanks to the huge interest and investments in the field of artificial intelligence (AI), this impact will immensely increase in the future. Together with the advances in the considerable computational power they need, DL algorithms can now seriously speed up some of the most demanding steps of the process and vastly expand the scale of the spaces to explore.^1–3^ The most striking example is the prediction of protein tridimensional structure by DL-based technologies. The remarkable performances of *eg*. AlphaFold^4^ or RoseTTAFold,^5^ together with the community-driven meticulous description of the applicability domain and biases, represent a transformative innovation in structural biology, bioinformatics and consequently influence hugely drug research.

Other areas of drug discovery have also benefited from DL but to a lesser extent. Drug design, *stricto sensu*, is one example. The inventing steps of crafting biologically relevant small molecules have always been very much human-driven. Medicinal and computational chemists are the molecular designers, and generally highly interdisciplinary experts. Medicinal chemistry has largely relied on Computer-Aided Drug Design (CADD) modeling for decades.^6,7^ Over 35 years ago, the first computational methods to conceive novel chemical structures from scratch - highly supervised to satisfy predefined molecular profiles, globally termed *de novo* design - were seen as a natural addition to the CADD arsenal.^8^ More recently, generative AI — predominantly DL techniques — were incorporated in *de novo* design. A paradigm shift was initiated, with drastically more automated and less supervised ways for the machine to build molecular structures. The vast number of virtual compounds generated by (deep) generative chemistry holds great promise with a few cases progressing to early-stage clinical development. However, to date no treatment exclusively designed by AI or DL has reached the market.^9^ It is unlikely that such a time will fully arrive in a near future, as the iterative trail- and-error cycles inherent to drug design will still rely on human decision-making based on a wealth of complex, diverse, and often intangible parameters.

Whether or not human expertise is involved, careful evaluation and prioritization of the AI-assisted generated chemical structures is imperative. Regrettably, only few studies describe generative chemistry for concrete medicinal chemistry applications.^10^ Even less are backed with experimental assessment (except recent outstanding exceptions^11,12^) and aim solely at optimizing molecules for simple computed parameters like *n*-octanol/water partition coefficient (log *P*), synthetic accessibility score (SA Score)^13^ or quantitative estimation of druglikeness (QED)^14^ for crudely evaluating descriptors of drug-like properties or ease of chemical synthesis. The origine of these arguable artificial properties are to be found in the first communications about automatic generation of one-dimensional SMILES (Simplified Molecular Input Line Entry System) strings^15^ to describe chemical structures. At that time, AI pioneers needed a gross estimation for the significance and relevance of these first machine-generated chemicals.^16–18^.

QED was proposed in 2012^14^ as an offshoot of the qualitative Lipinski’s Rule-of-five^19^ to quantify the drug-likeness nature by combining eight computed parameters (molecular weight, predicted Log *P*, number of H-bond donors, number of H-bond acceptors, topological polar surface area, number of rotatable bonds, number of aromatic rings and number of structural alerts^20^). Though many reported useful applications in conventional medicinal chemistry, distinction between drug and non-drug compounds was shown inaccurate,^21^ partly because of the too simplistic molecular and physicochemical descriptions but also because the range of properties of new drugs has broaden with time.^22^ Additionally, the QED was trained long before the emergence of generative chemistry and therefore the properties of (non-druglike) AI-crafted chemical structures cannot be captured properly making the differentiation even more difficult for such drug-likeness filter.^23^ Due to the straightforward calculation of a smooth, interpretable, continuous proxy of drug-likeness (between 0 and 1), QED is tail nowadays very frequently used as an evaluation metric for AI-generated compounds, even if it biases toward “average” accepted drug structures and disfavors novel chemotypes.^24^ For DL-generative methods like variational autoencoders (VAEs) or generative adversarial networks (GANs), QED can operate as an optimization target or assessment metric in order to lead the algorithm towards (too restricted) drug-likeness chemical space. For reinforcement learning, QED is routinely chosen as a reward function, to construct molecular structures alike known oral drugs. Optimization of drug-likeness alone is not of particularly interest in early phases of drug discovery, but QED represents one of the archetypal targets commonly chosen for multiple objective optimization strategies.^25^ Importantly, as mentioned above, QED was developed to assess whether molecules are alike actual drugs, and it predates the emergence of generative chemistry. It was therefore developed using a few molecules designed by medicinal chemists, which, by definition, were generally relevant to medicinal chemistry and passed all developments phases. As a result, QED is not suitable for determining if a molecule, beyond its drug-likeness, would be considered tractable, relevant, and interesting for drug discovery.

As already expressed, another ‘historical’ metric used to behold the relevance of virtual molecules is the synthetic accessibility score (SA Score)^13^, a rapid estimation of how difficult it would be to synthesize them. It is based on a fragmental contribution system built from large collections of synthesized compounds to quantify how frequently the fragments of a molecular structure to be evaluated appear in known molecules. The final score — from 1 for easy to 10 for difficult to synthetize — is obtained by summing fragmental contributions and penalties related to the size and the presence of complex moieties, like macrocycles, bridged ring systems or chiral centers. While SA Score concerns a separate property and is constructed differently, it shares similar limitations with QED when applied to the outputs of generative chemistry algorithms, for which they were not originally developed to provide guidance. Due to its additive nature, the SA Score tends to steer the generation process towards minimal, small molecules including few fragments. Also, because it is biased on data accentuating common fragments, it tends to favor trivial overly noncomplex, simplistic structures including usual chemical moieties.

A variety of benchmark schemes and validation sets have been proposed to quickly and reproducibly measure the performance of generative chemistry model. QED and SA Score are still ubiquitous in such widely adopted evaluation tools, either directly or indirectly. GuacaMol,^25^ a foundational benchmark for generative molecular models, uses QED directly as target objective functions but not SA Score. It has been shown however that it favors trivial, minimal chemical structures^26^. The renown benchmark suite MOSES^27^ evaluates the distribution of four properties described by molecular weight, log P, QED and SA Score as descriptive statistics for molecular sets, thus influencing the interpretation of model capacities. Validation procedures have been recently added to the DrugEx package,^28^ allowing users to select QED or SA Score as reward function parameters. This makes the process flexible, though it still largely relies on conventional metrics.

Both SA Score and QED suffer from selection bias, with at least 10-year-old training sets showing overrepresentation of certain chemotypes and lack of negative examples. Besides, they both lack a direct link with biological activity.^29^ Therefore, the combination of those two computational metrics, routinely employed as filters, reward functions, or constraints for generative chemistry incites such algorithms to find trivial ways of optimization and discourage exploration of more distant and broader chemical spaces, and ultimately to limits the innovative potential of AI-driven automatic chemical design.

Apart from employing more sophisticated pharmacokinetics models and retrosynthesis simulations when feasible — much more computationally demanding and difficult to implement at every step including the too simplistic scores — good practices for generative chemistry should implicate context-dependent expert interpretation and validation at key stages of the process. To address this need, we propose here an intuitive and highly interpretable tool — the OiiSTER-map for Optimal intuitively interpretable Structural Tool for Evaluating Real-world relevance—to characterize the relevance and significance of virtual molecules in the bioactive chemical space. This two-dimensional heatmap is built using molecular fingerprints and two key metrics, enabling a comprehensive visualization of structural complexity and relevance. The diverse analyses presented support the practicability of the evaluation tool, which offers a semi-qualitative classification of chemical structures and quantitative scoring of their usuality and degree of elaboration.

## Results

We characterized drug discovery-related molecules using two features. The first is the *number of bits*, defined as the number of bits set to 1 in its ECFP4 fingerprint.^30^ The second is the *usuality score*, calculated as the sum of frequencies of these bits across all molecules reported as active in binding assays in ChEMBL,^31^ considering only the 50% less common bits for the molecule. The first feature reflects the level of structural elaboration of the compound. Small values correspond to molecules with few distinct chemical substructures, as defined by ECFP4, while large values indicate molecules composed of many different moieties. Notably, *number of bits* does not necessarily correlate with molecular size. For example, octanoic, decanoic, and dodecanoic acids each have the same *number of bits* of 21, demonstrating that despite their size differences, their structural complexity is similar - they share the same chemical functions and differ only in the length of the carbon chain. The *usuality score* quantifies how common or rare the chemical features of a molecule are, based on their frequency in the ChEMBL dataset. A lower score indicates that a large portion of the molecule’s less frequent features are indeed very uncommon across other bioactive compounds. This suggests that the molecule may be chemically distinct from typical pharmacologycally relevant molecules—or, in some cases, even unrealistic or chemically invalid. In contrast, a high usuality score implies that the molecule consists mainly of standard, well-represented chemical features, in the medicinal chemistry context, making it likely chemically plausible, though potentially overly simplistic.

The *number of bits* and *usuality score* were calculated for the 675,279 molecules of the ChEMBL dataset, allowing to create a heatmap of their frequency at given values, called the OiiSTER-map. Results showed that bioactive molecules are not uniformely distributed on this heatmap, and that a large majority are found in a region roughly defined by 43 < *number of bits* < 73, and 0.2 < *usuality score* < 0.8. Based on this and on the visual inspection of molecules present in different locations of the (*number of bits* , *usuality score*) heatmap, several regions were defined. The right part of the heatmap containing molecules with more bits set to 1 in their fingerprint than the average was defined as containing the so-called “over-elaborated” compounds. On the contrary, the left part of the heatmap contains the so-called “minimal” molecule that contain substantially fewer chemical features than typical medicinal chemistry related molecules. In between, we find compounds that can be considered as “regular” in terms of number of different chemical features. The upper part of the heatmap is populated by molecules that can most of the time be considered as chemically “trivial” compared to typical drug discovery-related compounds. The opposite lower part contains molecules that are called “unusual”. In between can be found molecules considered as “Balanced” in term of *usuality score*. This decomposition of the OiiSTER-map into regions therefore allows to characterize a molecule, indicating if it well in the average of medicinal chemistry molecules in terms of chemical structure diversity and complexity, or if it is on the contrary too trivial, unusual, minimal or over-elaborated.

In the OiiSTER-map, most compounds gather within a region approximately defined by 42 < number of bits < 72 and 0.2 < usuality score < 0.8. Based on this distribution—and supported by visual inspection of randomly chosen molecules from various regions of the heatmap— distinct zones were defined (*Figure 1*):

**Figure 1.**
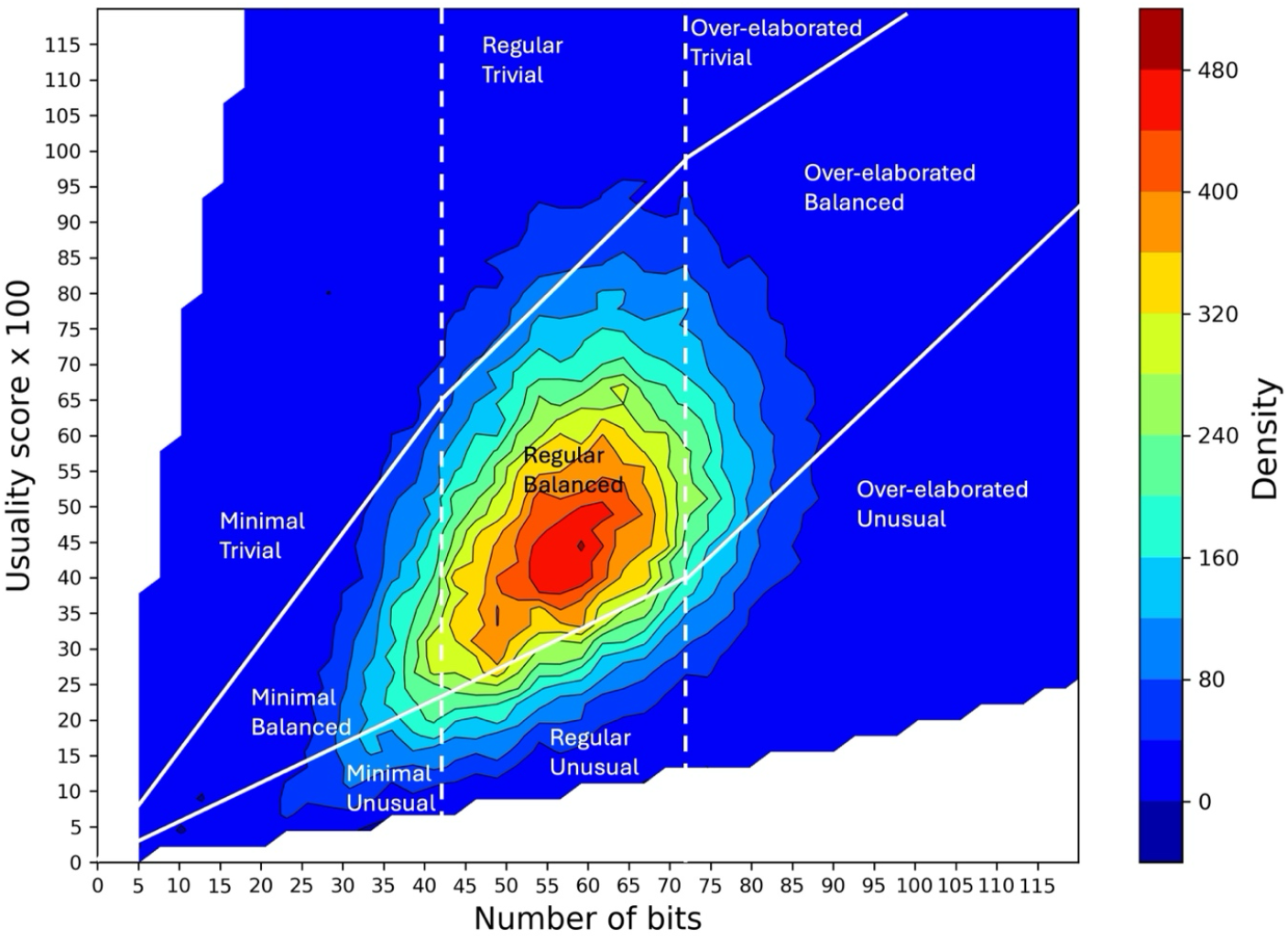
OiiSTER-map, heatmap of all molecules from the ChEMBL dataset as a function of their *number of bits* and *usuality score*. The color-coded density represents the number of molecules within each 1×1 bin used to discretize the 2D space into a regular grid prior to interpolation. Solid white lines divide the space into regions corresponding to trivial, balanced, and unusual molecules, while dotted white lines further partition it into zones of minimal, regular, and over-elaborated molecules. The full decomposition into 3×3 regions depicts nine classes in the complexity/usuality referential useful to qualify molecular structures.

- The right side of the heatmap, containing molecules with a higher-than-average number of bits, corresponds to “Over-elaborated” compounds, excessively rich in diverse chemical substructures.
- The left side, populated by molecules with substantially fewer features, represents “Minimal” compounds, which are structurally simpler than typical medicinal chemistry molecules.
- Molecules in the central zone in terms of *number of bits* fall within a typical range for both features and are referred to as “Regular”.
- The upper region includes molecules with very common and widely shared substructures, considered “Trivial” in comparison to typical drug-like compounds.
- In contrast, the lower region hosts molecules with rare or unconventional substructures, labeled “Unusual”.
- Between these extremes lies the “Balanced” region, comprising molecules with a moderate usuality score, reflecting a mix of common and less common features.

This heatmap thus provides a practical framework for characterizing molecules according to their structural complexity and chemical typicality. It helps determine whether a compound aligns with the core of bioactive chemical space or deviates toward being overly trivial, unusual, minimal, or over-elaborated.

Table 1 shows that 59.5 % of the molecules in the ChEMBL dataset fall within the Regular/Balanced class. In contrast, only about 12.4% are classified as Minimal and 14.2% as Over-elaborated, underscoring the predominance of Regular and Balanced compounds among real molecules synthesized within actual medicinal chemistry projects. On average, the molecules in the ChEMBL datase have a *number of bits* of 58 and a *usuability score* of 0.495 (Supplementary Material, Fig. S6).

**Table 1.**
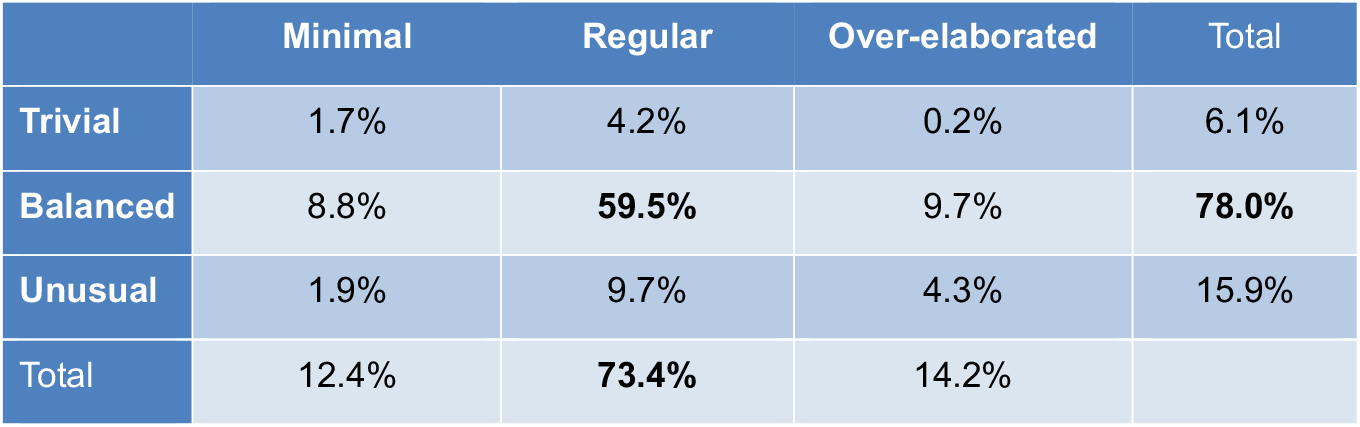
Fraction of bioactive molecules from the ChEMBL dataset belonging to the different classes of the OiiSTER-map.

As an initial test case, we assembled a set of molecules that are clearly too simplistic to be suitable for most medicinal chemistry applications (Supplementary Material File TestCompounds.xlsx). This set was complemented with compounds extracted from the main text or supplementary materials of various peer-reviewed publications presenting AI-based methods for *de novo* generation of drug discovery related molecules. From these sources, we deliberately selected a mix of compounds, some with clear medicinal chemistry relevance and others overly simplistic or chemically unrealistic, totalizing 99 compounds. As shown in Supplementary Material Table S1, our characterization approach successfully identified a substantial proportion of Minimal and Unusual compounds within this test set. Interesingly, of the 99 molecules analyzed, 63 passed conventional filtering criteria (QED ≥ 0.5 and SA Score ≤ 4). However, only 20 of these were classified as Regular and Balanced, while 40 were considered Minimal and 10 Unusual (see Supplementary Material File TestCompounds.xlsx). Figure 2 displays representative examples mapped onto the OiiSTER-map, demonstrating that even molecules meeting standard QED and SAscore ranges may not be sufficiently relevant for medicinal chemistry efforts. On the contrary, Supplementary Material Fig. S2 displays some molecule from the Regular/Balanced region that also passed the QED and SAscore filters. These last molecules can be considered relevant for drug discovery projects. This underscores the added value of our characterization framework in filtering AI-generated compounds beyond conventional metrics.

**Figure 2.**
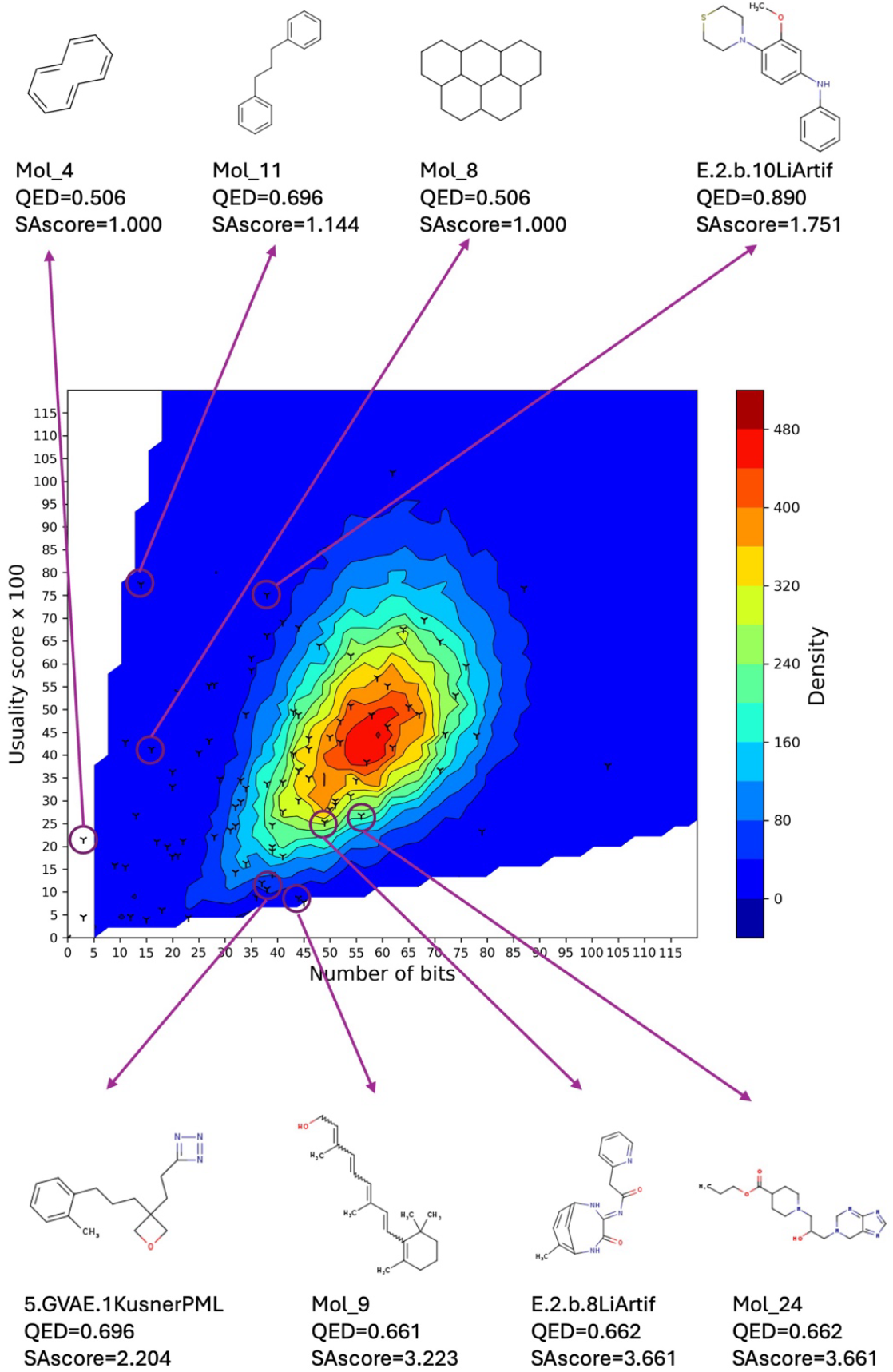
Location on the OiiSTER-map of the 99 molecules from the manually selected test set. Each cross corresponds to one of the 99 digital molecules. A few molecules that pass the QED and SA Score test, but are not in the Regular/Balanced region are examplified.

To demonstrate further the utility of the OiiSTER-map as a filtering tool for identifying irrelevant molecules possibly created by AI, we trained a LSTM model on 50,000 SMILES from the ChEMBL dataset. This model was deliberately sub-optimal to make the exercise more chalenging. This model was then used to generate 281 SMILES, referred to here as “digital molecules”. For each digital molecule, we computed QED, SA Score, *number of bits*, and *usuality score* (See Supplementary Material File LSTM_Compounds.xlsx). As shown in Table 2, while the Regular/Balanced class remains the most populated, the digital molecules include a higher proportion of Minimal and Unusual compounds compared to the ChEMBL dataset. This highlights the effectiveness of our classification system in revealing limitations of generative models. Interestingly, among the 133 digital molecules with a QED ≥ 0.5 and a SAscore ≤ 4—thresholds commonly used to indicate drug-like and synthetically accessible compounds—only 59 (44.4%) fall within the Regular/Balanced class of the characterization heatmap. Notably, all Over-elaborated molecules were excluded by these filters, and the proportion of Unusual compounds dropped from 39.4% to 18.0%. However, the fraction of Minimal compounds rose to 42.0%, largely because smaller molecules are often considered easy to synthesize while being able to meet drug-likeness criteria. Supplementary Material Figures S3, S4 and S5 display the location of these digital compounds, examplying molecules that pass the QED and SAcore filters, but fail or not to occupy the Regular/Balanced region. While the percentage of Regular/Balanced molecules increased after QED and Sascore filtering compared to the full set of 281 digital molecules, this result highlights again the limitations of relying solely on these two metrics. A substantial fraction of the filtered compounds still falls outside the optimal regions for medicinal chemistry. The OiiSTER-map thus offers a valuable complementary filter to refine AI-generated molecule selection and prioritize candidates better suited for drug discovery.

**Table 2.**
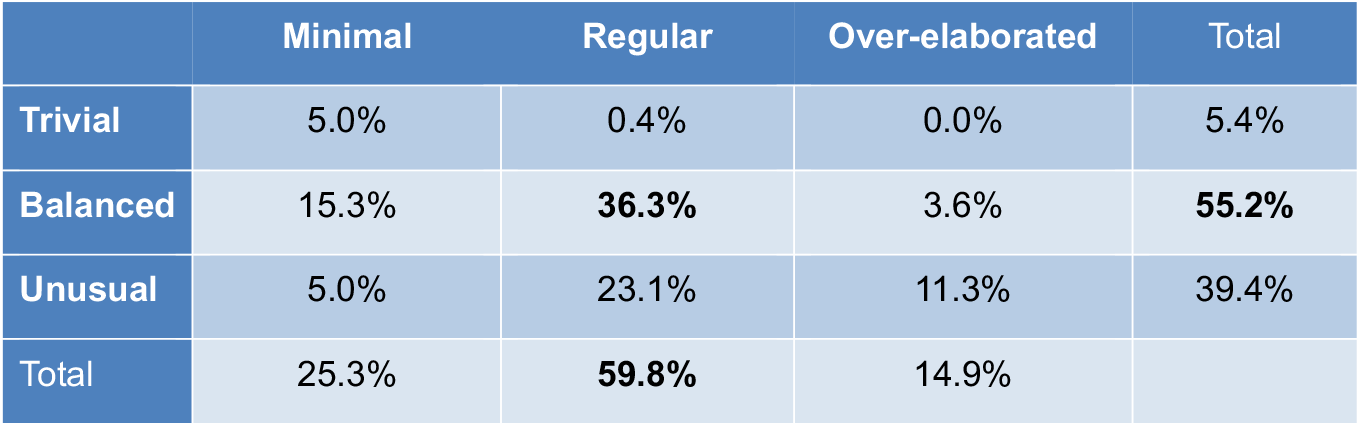
Fraction of the 281 digital molecules, generated by a simple sub-optimal LSTM deep-learning model, belonging to the different classes of the OiiSTER-map.

|  | Minimal | Regular | Over-elaborated | Total |
| --- | --- | --- | --- | --- |
| Trivial | 5.0% | 0.4% | 0.0% | 5.4% |
| Balanced | 15.3% | **36.3%** | 3.6% | **55.2%** |
| Unusual | 5.0% | 23.1% | 11.3% | 39.4% |
| Total | 25.3% | **59.8%** | 14.9% |  |

**Table 3.** Fraction of the 133 digital molecules, generated by a simple sub-optimal LSTM deep-learning model with QED ≥ 0.5 and SAscore ≤ 4, belonging to the different classes of the OiiSTER-map.

|  | Minimal | Regular | Over-elaborated | Total |
| --- | --- | --- | --- | --- |
| Trivial | 8.2% | 0.8% | 0.0% | 9.0% |
| Balanced | 28.6% | **44.4%** | 0.0% | **73.0%** |
| Unusual | 5.2% | 12.8% | 0.0% | 18.0% |
| Total | 42.0% | **58.0%** | 0.0% |  |

Interestingly, the information in the heatmap is directly related to bioactivity. Actives with low experimental affinity for their targets, as annotated in the ChEMBL dataset, tend to be more Minimal than the dataset average, although they still fall within the Regular/Balanced class on average. In contrast, high-affinity compounds are generally more Elaborated, yet their average *number of bits* and *usability scores* also remain within the Regular/Balanced region (Supplementary Material, Fig. S6). This trend likely reflects the fact that achieving higher affinity often requires an increased number of complementary interactions between the ligand and the target protein, which in turn demands a larger molecular size and a richer array of chemical functionalities.

Based on the dates of reaching clinical phase 4, as annotated in ChEMBL, we analyzed the distribution of approved drugs on the OiiSTER-map. Interestingly, drugs approved since 1900 span a broad area of the heatmap, extending well beyond the Regular/Balanced region— particularly toward the left, where Minimal compounds are found (Supplementary Material, Fig. S7 top). A deeper analysis revealed a marked shift in the nature of approved drugs over time (Supplementary Material, Fig. S7 and S8). Compounds approved between 1900 and 2000 tend to be significantly more Minimal than the average ChEMBL dataset molecules. During this period, the Minimal/Balanced region was the most populated (30.2% of approved drugs), followed closely by the Regular/Balanced region (27.8%). Among the Minimal pre-2000 approved drugs, we can typically find amphetamine, acetaminophen, ibuprofen, diclofenac or sulfanilamide, as examples. In contrast, drugs approved more recently—from 2010 to 2024— display a distribution *of number of bits* and *usability scores* closely aligned with the overall ChEMBL dataset. In this group, the Regular/Balanced region dominates, encompassing 50.0% of the approved drugs, while only 8.5% fall within the Minimal/Balanced space. This trend likely reflects a growing emphasis on developing potent and selective ligands, which often requires larger molecules with more diverse and functionally rich chemical structures. Notably, in recent years, a significant fraction of approved drugs—ranging from 15% to 19% since 2000—are classified as Over-elaborated. These include large natural or semi-synthetic compounds such as telithromycin and everolimus, polypeptides like atosiban, and sizable synthetic molecules like venetoclax. Notably, telithromycin and everolimus are classified as particularly Unusual, with *usuality scores* of 0.25 and 0.29, respectively, and a *number of bits* of 111 and 122. This aligns with their origin as derivatives of natural compounds, in contrast to most molecules in the ChEMBL dataset. Conversely, venetoclax is categorized as Balanced, with 114 bits and a high *usuality score* of 0.96. Atosiban, with 115 bits and a *usuality score* of 0.55, is also considered Unusual—though to a lesser extent than telithromycin and everolimus.

We also analyzed compounds that reached Phase 1 (353 molecules), Phase 2 (1,451 molecules), or Phase 3 (453 molecules), irrespective of whether they were discontinued at that particular stage or are still under investigation. As shown in Supplementary Figure S9, the distribution of Phase 1 molecules across the OiiSTER-map closely mirrors that of the overall ChEMBL dataset. However, a noticeable increase in the proportion of Minimal compounds is observed among those that reached up to Phases 2 and 3. One possible explanation is that Minimal molecules may generally exhibit lower binding affinity and selectivity for their targets— as previously discussed and shown in Supplementary Figure S6—which could lead to discontinuation in those phases due to insufficient efficacy.

### General Discussion

The OiiSTER-map, proposed here as a molecular structure evaluation tool, introduces a qualitative and quantitative framework that shows clear complementarity to and offers arguable advantages over traditional metrics, especially for evaluating virtual molecules generated by AI models in drug discovery.

Most employed traditional evaluation metrics, QED and SA Score, are mainly based on approved drugs and on organic synthetic compounds, respectively, lacking straightforward relationship with biological relevance.^29^ In contrast, the OiiSTER-map is directly linked to highly curated bioactivity knowledge from ChEMBL through the density quantification and the *usuality score*, and hence emphasizes pharmacologically relevant structural features and not just chemically plausible ones (refer to Supplementary Fig. S6). Besides, one strong point is the decomposition of the molecular description along two orthogonal axis that allows for more detailed insights on why a generated chemical structure might have difficulties to reach stepwise drug discovery objectives, with human-interpretable criteria like “too minimal or over-elaborated molecule” or “trivial or implausible in bioactivity space”. The 3×3 regional decomposition over the usuality *vs*. number of bits heatmap (a proxy of complexity or level of elaboration) provides nuanced evaluation categories (see Fig. 1). We can anticipate that in most applications, a human or a machine will favor a “Balanced/Regular” classification or filter out what is distant from this drug discovery chemical “sweet spot”. However, the OiiSTER-map allows medicinal chemists to appreciate other promising spaces that can help to balance innovation, discovery and feasibility. For instance, the identification of “Unusual/Minimal” region could contain potentially overlooked innovative but simple compounds. On the contrary, an expert could search into “Trivial/Over-elaborated” class for problematic extreme compounds but showing established bioactive patterns (*eg*. pharmacophores) that may be simplified through chemical optimization to discover novel chemotypes. This exploration capacities in the referential of the OiiSTER-map can also be leveraged for comparative or benchmark activities. The outputs of various generative models—or different parameter settings of a single model—can be visualized by mapping the chemical space occupied by the resulting molecular structures directly onto the heatmap.

Those qualitative human-in-the-loop decision-making analyses are not the only applications foreseen for the OiiSTER-map. The descriptors being numerical values, both scores are suitable for implementation in AI-assisted drug design methods. Both scores integrated inside generative loops DL algorithms (e.g. GANs or VAEs) as optimization targets or assessment parameters could efficiently steer the process toward a user-defined space on the heatmap (in most cases probably for “regular” complexity and “balanced” usuality). Likewise in reinforcement learning both scores can be chosen as reward functions to craft molecular structures of desire properties in the bioactivity complexity/usuality referential. Also, they could enable multi-objective optimization with better interpretability. The practical feasibility and impact of such integration in generative chemistry, combined with other possibly complementary metrics, is certainly one of major perspectives of this preliminary work. Possible future developments of the method can be foreseen: ChEMBL provides standardized, highly curated bioactivity source, but also gives access to numerous additional information regarding the biological or the chemical contexts. This opens the way to possible indication-specific or target-specific heatmaps by filtering data. Also, additional weighting parameters like ranges of potency for the active compounds or target classes for the proteins could help refine the description of the usuality/complexity chemical space on the OiiSTER-map.

Based on the long experience on the heavy usage of our BOILED-Egg model for ADME,^32^ we anticipate that the here-proposed OiiSTER-map will benefit from its conceptual simplicity, speed and user-friendliness. Indeed, the calculation of ECFP4 molecular fingerprints, the count of bits and the comparison with reference binaries are extremely fast and scalable. This makes the tool suited for implementation either in custom-made generative chemistry models for guidance or in web applications like SwissADME^33^ for real-time evaluation of generated structure.

We foresee that a routine usage of a bioactivity-informed, structurally aware, and easily interpretable tool like the OiiSTER-map, in combination with other well implemented metrics, can be decisive to go beyond current limitations of assessing (deep) generative models and to ensure a more mechanistically relevant, nuanced and useful evaluation of the thus designed virtual molecules.

## Methods

### Data extraction and ECFP4 fingerprints generation

Compounds were extracted from ChEMBL^31^ (version 34) using a MySQL query that retrieved all molecules in SMILES format^15^ with a molecular weight lower than 1000 g/mol, tested in binding assays (assay type “B”, confidence score >3) on macromolecular targets (single proteins or protein complexes), and with IC_50_, K_D_ or K_i_ values <1 mM. This yielded 675,908 unique SMILES. The structures were standardized using JChem Microservices (v21.3, www.chemaxon.com) through desolvation, unsalting, neutralization, and kekulization. Each standardized SMILES was then converted to an Extended Connectivity Fingerprint (ECFP4), encoded in hexadecimal format, using OpenBabel (v3.1.1).^34^ These were subsequently transformed into 4096-bit binary vectors, each representing the presence or absence of specific circular substructures. Standardization and fingerprinting yielded ECFP4 binary vectors^30^ for 675,279 molecules, each identified by its ChEMBL ID, and called the ChEMBL dataset below.

For each molecule, the ChEMBL IDs and the affinity for each target were retained for downstream analysis. When the same molecule was tested several times, the best affinity it achieved was retained. When available in ChEMBL, the maximum clinical trial phase reached by each molecule (ranging from Phase 1 to Phase 4) was recorded, along with the year it was reached. Phase 4 indicates regulatory approval and market release.

### Calculation of the OiiSTER-map scores

The OiiSTER-map we introduce here to characterize quantitatively molecules is based on two scores, the *number of bits* and the *usuality score* (Supplementary Material Fig. S1).

The first one, used for the x-axis, is simply the number of bits set to 1 in the molecule’s ECFP4 fingerprint, computed by summing the 1s in the binary vector.

The *usuality score*, used for the y-axis, is computed in two steps. First, the frequency with which a given bit was set to 1 in the ECFP4 fingerprints of all molecules in the ChEMBL dataset were calculated for each bit. These frequencies range from 0 for the less frequent circular substructures to 1 for the most frequent ones. The frequencies are precomputed once for the entire dataset. Then, for a given molecule, the bits set to 1 in its ECFP4 fingerprint are ranked by their frequency. The *usuality score* is calculated as the sum of the frequencies of the 50% less common bits (Supp Mat Fig. S1).

Heatmaps were generated using Matplotlib (v3.6.3) with cubic interpolation. The interpolation was applied to a grid file containing the number of ChEMBL molecules in each 1×1 bin defined by *number of bits* and *usuality score* x 100.

### Chemoinformatics

Molecular images were generated using the JChem Microservices (version 21.3, www.chemaxon.com). The Quantitative Estimate of Drug-likeness (QED)^14^ values and the SA Score^13^ were calculated using RDKIT, version 202309.1.

### Simple DL generative chemistry model

A simple Deep Learning (DL) model was used to generate small molecules. SMILES of 50,000 randomly chosen molecules from the ChEMBL dataset were used to train a Long Short Term Memory (LSTM) model using Tensorflow (version 2.19). Before training, the SMILES were concatenated, separating molecules by a dot. The model was trained during 20 epochs, using a batch size of 128, the RMSprop Keras optimizer, a learning rate of 0.01 and using categorical cross entropy as loss function. At the end of the training a character chain of 40,000 characters was generated by the model, and treated as a list of SMILES separated by dots. This led to a total of 605 SMILES, of which 281 unique SMILES were readable by RDKIT and considered as putative molecules. These SMILES contained a range of 8 to 123 characters. QED, SAsc ore, *number of bits* and *usuality score* were calculated for each of them.

## Supporting information

The 99 test compounds

The 281 molecules generated using a simple LSTM model

## Author contributions

V.Z. conceptualized and supervised. A.D. and V.Z. scripted, performed analyses, and wrote, reviewed and accepted the manuscript.

## Acknowledgement

We acknowledge ChemAxon (www.chemaxon.com) for the licensing agreement.

## References

1. Tropsha, A.Martin, H.-J. & Cherkasov, A. The Six Ds of Exponentials and drug discovery: A path toward reversing Eroom’s law. Drug Discov. Today 30, 104341 (2025).

2. Gangwal, A. & Lavecchia, A. Unleashing the power of generative AI in drug discovery. Drug Discov. Today 29, 103992 (2024).

3. Wang, H. et al. Scientific discovery in the age of artificial intelligence. Nature 620, 47–60 (2023).

4. Abramson, J. et al. Accurate structure prediction of biomolecular interactions with AlphaFold 3. Nature 630, 493–500 (2024).

5. Lisanza, S. L. et al. Multistate and functional protein design using RoseTTAFold sequence space diffusion. Nat. Biotechnol. 1–11 (2024) doi:10.1038/s41587-024-02395-w.

6. Pitt, W. R. et al. Real-World Applications and Experiences of AI/ML Deployment for Drug Discovery. J. Med. Chem. 68, 851–859 (2025).

7. Hasselgren, C. & Oprea, T. I. Artificial Intelligence for Drug Discovery: Are We There Yet? Annu. Rev. Pharmacol. Toxicol. 64, 527–550 (2023).

8. Schneider, G. & Fechner, U. Computer-based de novo design of drug-like molecules. Nat. Rev. Drug Discov. 4, 649–663 (2005).

9. Jayatunga, M. K., Ayers, M., Bruens, L., Jayanth, D. & Meier, C. How successful are AI-discovered drugs in clinical trials? A first analysis and emerging lessons. Drug Discov. Today 29, 104009 (2024).

10. Bilodeau, C., Jin, W., Jaakkola, T., Barzilay, R. & Jensen, K. F. Generative models for molecular discovery: Recent advances and challenges. Wiley Interdiscip. Rev.: Comput. Mol. Sci. 12, (2022).

11. Atz, K. et al. Prospective de novo drug design with deep interactome learning. Nat. Commun. 15, 3408 (2024).

12. Chen, S. et al. Deep lead optimization enveloped in protein pocket and its application in designing potent and selective ligands targeting LTK protein. Nat. Mach. Intell. 7, 448–458 (2025).

13. Ertl, P. & Schuffenhauer, A. Estimation of synthetic accessibility score of drug-like molecules based on molecular complexity and fragment contributions. J. Cheminform. 1, 8 (2009).

14. Bickerton, G. R., Paolini, G. V., Besnard, J., Muresan, S. & Hopkins, A. L. Quantifying the chemical beauty of drugs. Nature Chem 4, 90–98 (2012).

15. Weininger, D. SMILES, a chemical language and information system. 1. Introduction to methodology and encoding rules. J. Chem. Inf. Comput. Sci. 28, 31–36 (1988).

16. Jaques, N. et al. Sequence Tutor: Conservative Fine-Tuning of Sequence Generation Models with KL-control. arXiv (2016) doi:10.48550/arxiv.1611.02796.

17. Kusner, M. J., Paige, B. & Hernández-Lobato, J. M. Grammar Variational Autoencoder. arXiv (2017) doi:10.48550/arxiv.1703.01925.

18. Gómez-Bombarelli, R. et al. Automatic Chemical Design Using a Data-Driven Continuous Representation of Molecules. ACS Cent. Sci. 4, 268–276 (2018).

19. Lipinski, C. A., Lombardo, F., Dominy, B. W. & Feeney, P. J. Experimental and computational approaches to estimate solubility and permeability in drug discovery and development settings. Adv Drug Deliv Rev 46, 3–26 (2001).

20. Brenk, R. et al. Lessons learnt from assembling screening libraries for drug discovery for neglected diseases. ChemMedChem 3, 435–444 (2008).

21. Beker, W., Wołos, A., Szymkuć, S. & Grzybowski, B. A. Minimal-uncertainty prediction of general drug-likeness based on Bayesian neural networks. Nat. Mach. Intell. 2, 457–465 (2020).

22. Shultz, M. D. Two Decades under the Influence of the Rule of Five and the Changing Properties of Approved Oral Drugs. J. Med. Chem. 62, 1701–1714 (2019).

23. Lee, K., Jang, J., Seo, S., Lim, J. & Kim, W. Y. Drug-likeness scoring based on unsupervised learning. Chem. Sci. 13, 554–565 (2021).

24. Bilodeau, C., Jin, W., Jaakkola, T., Barzilay, R. & Jensen, K. F. Generative models for molecular discovery: Recent advances and challenges. Wiley Interdiscip. Rev.: Comput. Mol. Sci. 12, (2022).

25. Brown, N., Fiscato, M., Segler, M. H. S. & Vaucher, A. C. GuacaMol: Benchmarking Models for de Novo Molecular Design. J. Chem. Inf. Model. 59, 1096–1108 (2019).

26. Shimizu, Y. et al. AI-driven molecular generation of not-patented pharmaceutical compounds using world open patent data. J. Cheminformatics 15, 120 (2023).

27. Polykovskiy, D. et al. Molecular Sets (MOSES): A Benchmarking Platform for Molecular Generation Models. Front. Pharmacol. 11, 565644 (2020).

28. Šícho, M. et al. DrugEx: Deep Learning Models and Tools for Exploration of Drug-Like Chemical Space. J. Chem. Inf. Model. 63, 3629–3636 (2023).

29. Liu, X. et al. MolFilterGAN: a progressively augmented generative adversarial network for triaging AI-designed molecules. J. Cheminformatics 15, 42 (2023).

30. Rogers, D. & Hahn, M. Extended-Connectivity Fingerprints. J. Chem. Inf. Model. 50, 742–754 (2010).

31. Zdrazil, B. et al. The ChEMBL Database in 2023: a drug discovery platform spanning multiple bioactivity data types and time periods. Nucleic Acids Res. 52, D1180–D1192 (2023).

32. Daina, A. & Zoete, V. A BOILED-Egg To Predict Gastrointestinal Absorption and Brain Penetration of Small Molecules. ChemMedChem 11, 1117–1121 (2016).

33. Daina, A., Michielin, O. & Zoete, V. SwissADME: a free web tool to evaluate pharmacokinetics, drug-likeness and medicinal chemistry friendliness of small molecules. Sci. Rep. 7, 42717 (2017).

34. O’Boyle, N. M. et al. OpenBabel: An open chemical toolbox. J. Cheminform. 3, 33 (2011).

