## Supplementary Figures for "Rethinking Molecular Beauty in the Deep Learning Era"

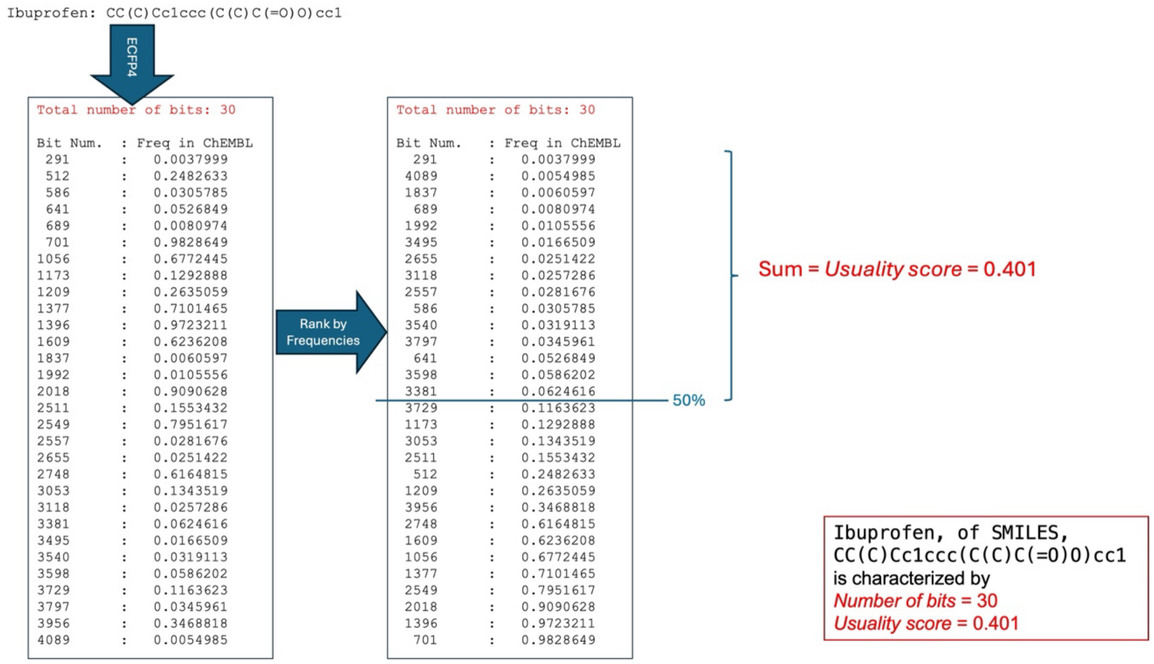


**Figure S1. Example of *number of bits* and *usuality score* calculation for the ibuprofen molecule.** First, the 4096-bit ECFP4 fingerprint of ibuprofen is generated using OpenBabel. Thirty bits are set to 1 in it, yielding a *number of bits* of 30. These bits are then ranked by their frequency across the ChEMBL dataset. The *usuality score* is computed as the sum of the frequencies for the 50% less common bits. For ibuprofen, the usuality score is 0.401.

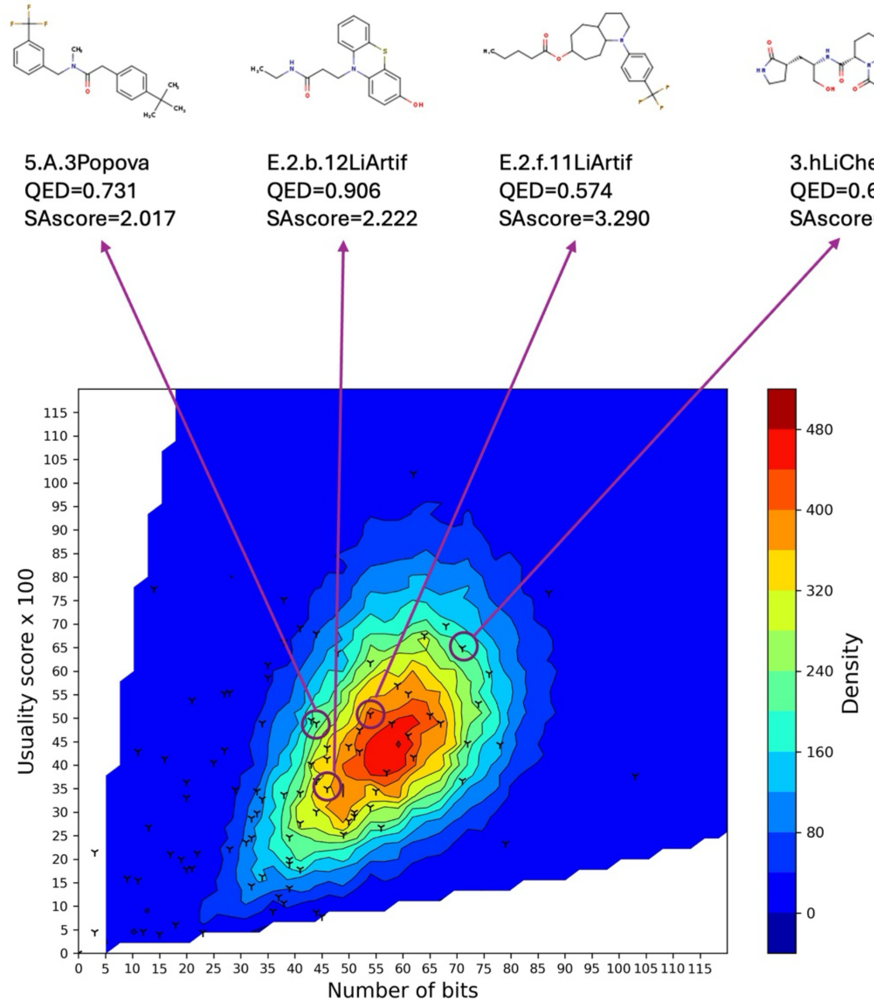


**Figure S2. Location on the characterization heatmap of the 99 molecules from the manually selected test set.** Each cross corresponds to one of the 99 test molecules. A few molecules that pass the QED and SAscore test, and are in the Regular/Balanced region are examplified.

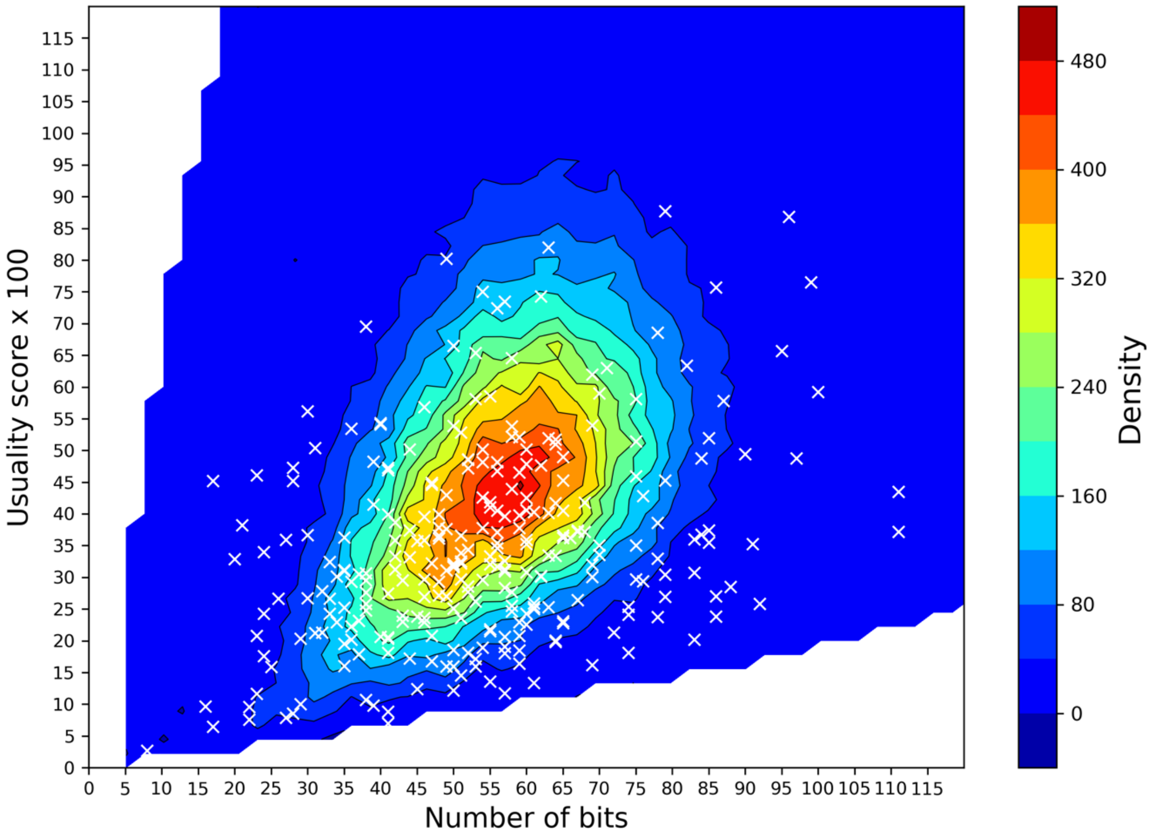


**Figure S3. Location of the 281 molecules generated by LSTM on the characterization heatmap.** Each cross corresponds to one of the 281 digital molecules.

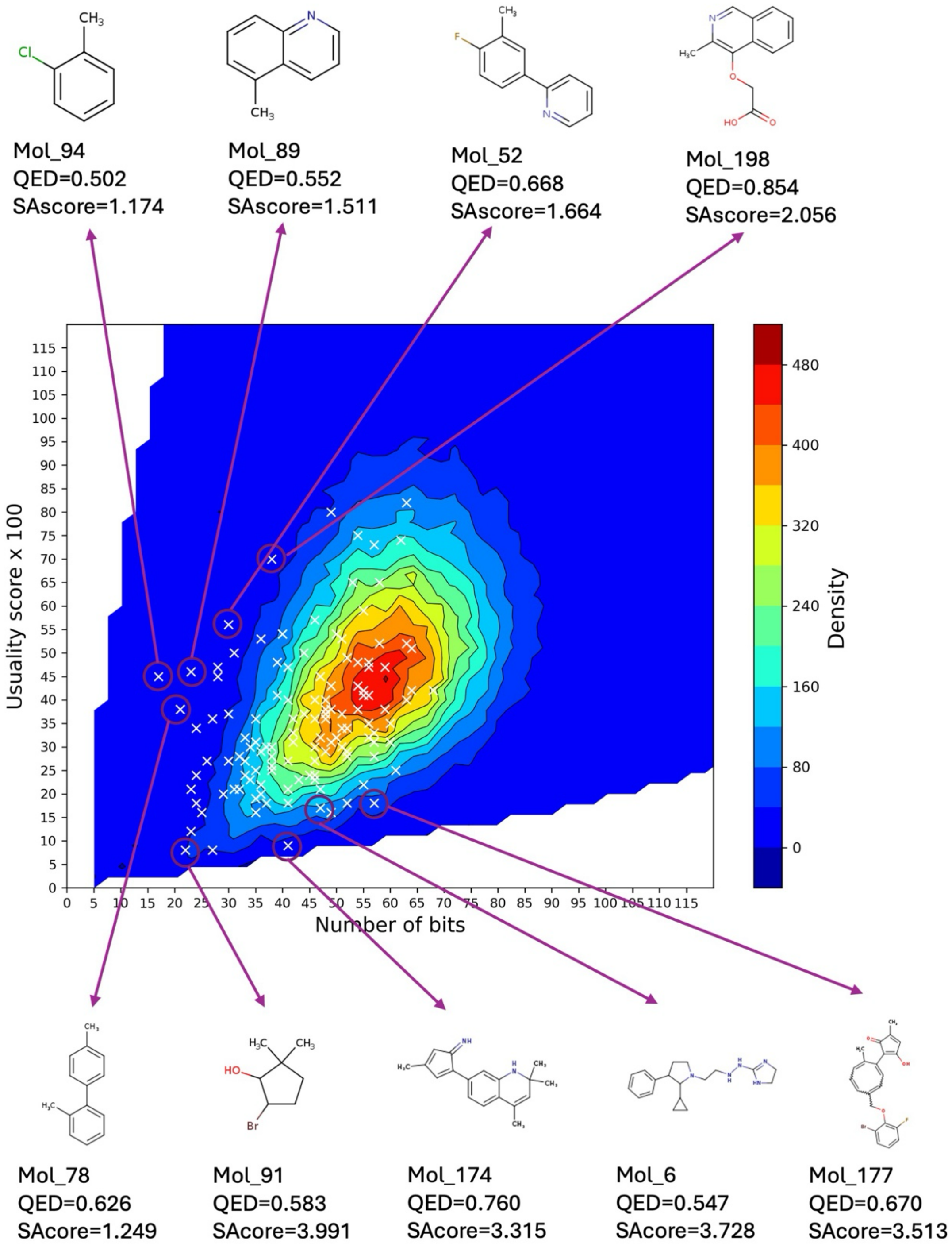


**Figure S4. Location on the characterization heatmap of the 133 molecules generated by LSTM that pass the QED ≥ 0.5 and SAscore ≤ 4 filters.** Each cross corresponds to one of the 133 digital molecules. A few examples of digital molecules that are obviously over-simplistic or chemically problematic are given.

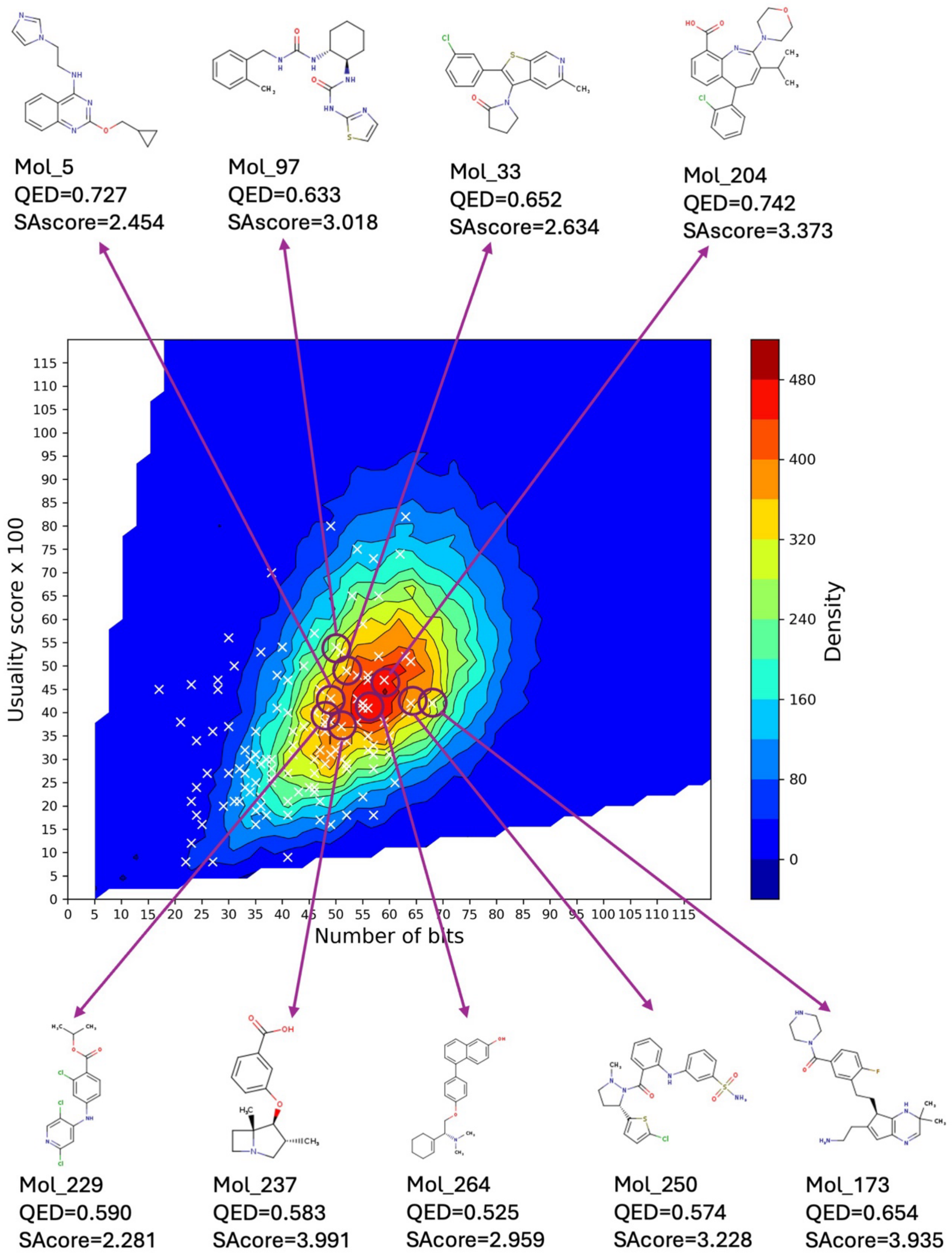


**Figure S5. Position on the characterization heatmap of the 133 molecules generated by LSTM that pass the QED ≥ 0.5 and SAscore ≤ 4 filters.** Each cross corresponds to one of the 133 digital molecules. A few examples of digital molecules that belong to the Regular/Balanced region are given.

| | | # Compounds | *Num. Bits* | *Usuality Score* |
|---|---|---|---|---|
| | All ChEMBL dataset | 675,279 | 57.7 | 0.495 |
| 1 | 1’000’000 nM ≥ Affinity > 100’000 nM | 10,916 | 46.9 | 0.388 |
| 2 | 100’000 nM ≥ Affinity > 10’000 nM | 53,932 | 50.8 | 0.435 |
| 3 | 10’000 nM ≥ Affinity > 1000 nM | 107,868 | 53.3 | 0.472 |
| 4 | 1000 nM ≥ Affinity > 100 nM | 151,619 | 56.4 | 0.496 |
| 5 | 100 nM ≥ Affinity > 10 nM | 173,058 | 59.3 | 0.511 |
| 6 | 10 nM ≥ Affinity > 1 nM | 124,361 | 62.0 | 0.515 |
| 7 | 1 nM ≥ Affinity > 0.1 nM (#44’314) | 44,314 | 64.7 | 0.524 |
| 8 | 0.1 nM ≥ Affinity | 9242 | 67.3 | 0.520 |

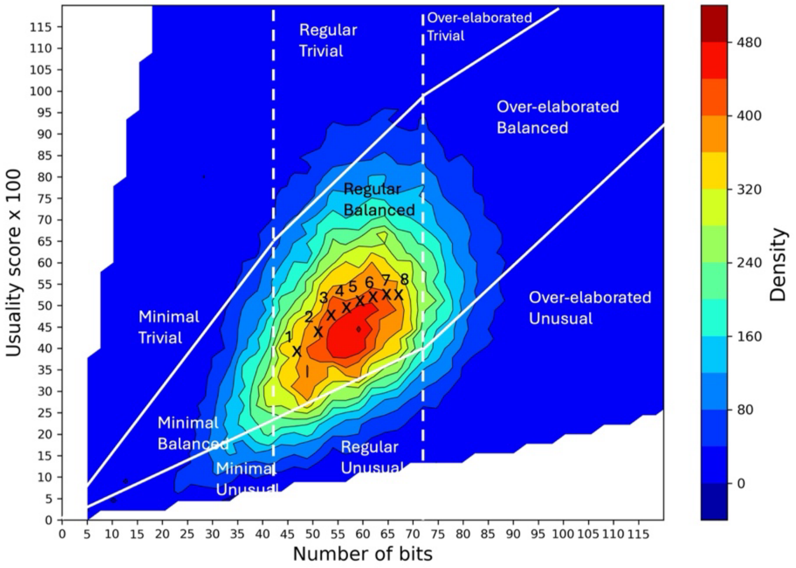


**Figure S6. Evolution of the average *number of bits* and *usuality score* of compounds from the ChEMBL dataset, as a function of their affinity for their target.** Crosses numbered 1 to 8 in the characterization heatmap correspond to the different affinity levels on the left table. The table reports the number of compounds achieving a given range of affinity (# Compounds) as well as their average *number of bits* and *usuality score*.

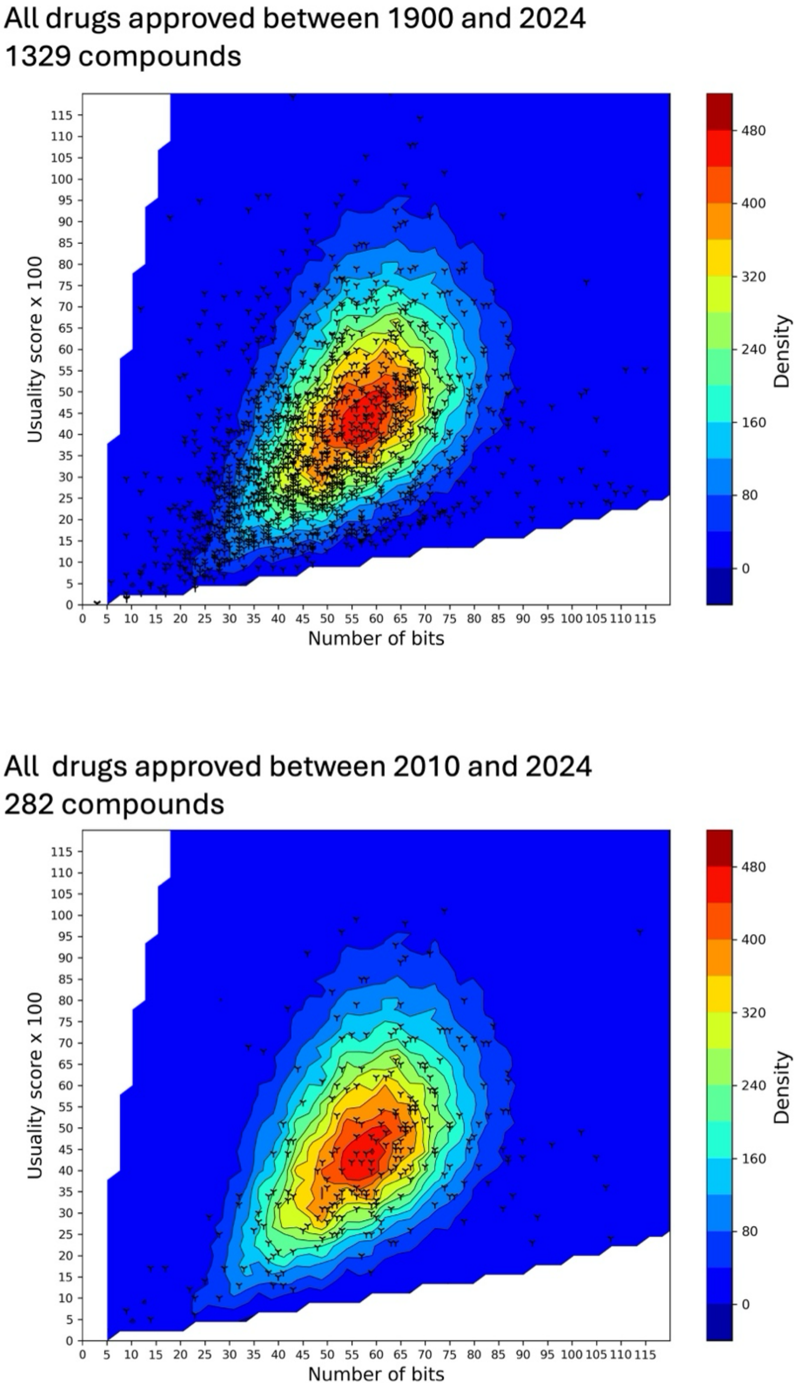


**Figure S7. Evolution of the position of phase 4, marketed drugs, on the characterization heatmap according to approval period.** Each cross corresponds to an approved drug. Top panel: Drugs approved by the FDA between 1900 and 2024 (n=1329); Bottom panel: Drug approved by the FDA between 2010 and 2024 (n=282)

Phase 4 drugs, approved in [1900, 2000[ (#851)

| | Minimal | Regular | Over-elaborated | Total |
|---|---|---|---|---|
| Trivial | 6.2% | 1.3% | 0.0% | 7.5% |
| Balanced | **30.2%** | 27.8% | 1.8% | **59.8%** |
| Unusual | 8.8% | 19.3% | 4.6% | 32.7% |
| Total | 45.2% | **48.4%** | 6.4% | |

Phase 4 drugs, approved in [2000, 2024[ (#478)

| | Minimal | Regular | Over-elaborated | Total |
|---|---|---|---|---|
| Trivial | 3.1% | 4.0% | 0.2% | 7.3% |
| Balanced | 11.7% | **49.2%** | 6.5% | **67.4%** |
| Unusual | 4.6% | 12.6% | 8.1% | 25.3% |
| Total | 19.4% | **65.8%** | 14.8% | |

Phase 4 drugs, approved in [2010, 2024[ (#282)

| | Minimal | Regular | Over-elaborated | Total |
|---|---|---|---|---|
| Trivial | 2.5% | 4.3% | 0.3% | 7.1% |
| Balanced | 8.5% | **50.0%** | 8.2% | **66.7%** |
| Unusual | 5.0% | 10.6% | 10.6% | 26.2% |
| Total | 16.0% | **64.9%** | 19.1% | |

Drugs become more elaborated over time

**Figure S8. Evolution of the characteristics of phase 4, marketed, drugs as a function of their approval date.** Tables show the fraction of approved drugs belonging to the different classes of the characterization heatmap, according to three ranges of approval dates, i.e. [1900, 2000[, [2000, 2024[ and [2010, 2024[, for a total of 851, 478 and 282 molecules, respectively.

Phase 1 drugs (#353)

| | Minimal | Regular | Over-elaborated | Total |
|---|---|---|---|---|
| Trivial | 3.1% | 2.5% | 0.0% | 5.6% |
| Balanced | 8.2% | **56.7%** | 9.1% | **74.0%** |
| Unusual | 2.3% | 10.2% | 7.9% | 20.4% |
| Total | 13.6% | **69.4%** | 17% | |

Phase 2 drugs (#1451)

| | Minimal | Regular | Over-elaborated | Total |
|---|---|---|---|---|
| Trivial | 4.3% | 2.5% | 0.0% | 6.8% |
| Balanced | 17.0% | **49.0%** | 6.5% | **72.5%** |
| Unusual | 4.4% | 12.8% | 3.5% | 20.7% |
| Total | 25.7% | **64.3%** | 10.0% | |

Phase 3 drugs (#453)

| | Minimal | Regular | Over-elaborated | Total |
|---|---|---|---|---|
| Trivial | 4.0% | 2.6% | 0.4% | 7.0% |
| Balanced | 15.2% | **41.8%** | 9.6% | **66.6%** |
| Unusual | 6.4% | 14.3% | 5.7% | 26.4% |
| Total | 25.6% | **58.7%** | 15.7% | |

**Figure S9. Evolution of the characteristics of candidate drugs as a function of their maximal clinical phase.** Tables show the fraction of drug-candidates in development belonging to the different classes of the characterization heatmap. Phase 1: 453 molecules; Phase 2: 1451 molecules; Phase 3: 453 molecules.

| | **Minimal** | **Regular** | **Over-elaborated** | Total |
|---|---|---|---|---|
| **Trivial** | 25.3% | 3.0% | 0.0% | 28.3% |
| **Balanced** | 18.2% | 31.3% | 3.0% | 52.5% |
| **Unusual** | 10.1% | 6.1% | 3.0% | 19.2% |
| Total | 53.6% | 40.4% | 6.0% | |

*Table S1. Fraction of molecules from the testset located in the different classes of the characterization heatmap.*

**Supplementary file TestCompounds.xlsx:** contains the ID, SMILES notation, calculated QED, SA Score, number of bits, usuality score and class on the OiSTER map for the 99 test compounds.

**Supplementary file LSTM_Compounds.xlsx:** contains the ID, SMILES notation, calculated QED, SA Score, number of bits, usuality score and class on the OiSTER map for the 281 molecules generated using a simple LSTM model.
